# Revealing α oscillatory activity using voltage-sensitive dye imaging in monkey V1

**DOI:** 10.1101/810325

**Authors:** Sandrine Chemla, Sebastien Roux, Alexandre Reynaud, Frédéric Chavane, Rufin VanRullen

## Abstract

The relevance of α oscillations (7-12Hz) in neural processing, although recognized long ago, remains a major research question in the field. While intensively studied in humans, α oscillations appear much less often investigated (and observed) in monkeys. Here we wish to provide data from non-human primates on stimulus-related α rhythm. Indeed, in humans, EEG α is enhanced in response to non-periodic dynamic visual stimulation (“perceptual echoes” or to a static stimulus (“flickering wheel illusion”). Do the same visual patterns induce an oscillatory response in monkey V1? We record voltage-sensitive dye signals from three anesthetized monkeys to investigate the population-based oscillatory neural response that is not resulting from attention-related feedback signals. We revealed α oscillations in monkey V1 which, when they occur, react in a manner comparable to human studies.

## Introduction

The occipital α rhythm is the most dominant rhythm in the awake human brain. It has been known since Hans Berger’s work in 1929 (Berger, 1929) that when you relax with eyes closed, large amplitude rhythmic oscillations between 7-12 Hz appear above the visual cortex. Thereby, studies of the EEG α rhythm in humans abound in the literature. Yet its functional role and relation to visual perception are still little understood and remain a major subject of research (Thut et al., 2006; Hanslmayr et al., 2007; Busch et al., 2009; Jensen and Mazaheri, 2010; Lange et al., 2014; Milton et al., 2016). In comparison, α oscillations have surprisingly not received much attention in monkeys’ physiological research (Jensen et al., 2015), compared to other brain rhythms (Eckhorn et al., 1993; Schmiedt et al., 2014). This discrepancy could come from the fact that α is generally suppressed by, or negatively correlated with visual stimulation (Klimesch et al., 2007), so this rhythm has long been viewed as the “idling” state of the brain (Pfurtscheller et al., 1996). However, α is actually not only a spontaneous oscillation: under certain conditions, it can also be positively driven by the visual stimulus, in relation to perception. In this paper, we focused on this stimulus-related α, i.e. increase in α amplitude evoked (or induced; David, Kilner & Friston, 2006) by the stimulus.

First, EEG α in humans is enhanced in response to random non-periodic dynamic stimulation (“perceptual echoes”, VanRullen and Macdonald, 2012). In that experiment, EEG was recorded while participants watched random dynamic sequences of luminance values (“white noise”, with a flat power spectrum) within a peripheral disc stimulus. The cross-correlation between the stimulus luminance sequence on each trial and the corresponding EEG was used to probe the impulse response of the brain’s visual system. After early modulations (at lags below 250ms) reflecting the classic visual evoked potential response, a striking 10 Hz oscillation was visible, as if the brain echoed the stimulation sequence for more than 1 second. In comparison, the cross-correlation between the same stimulus sequence and a randomly chosen EEG trial did not show the echo. The topographical mapping of this 10 Hz echo in the cross-correlation functions revealed that the oscillation principally occurred over occipital electrodes, which suggests a role for the occipital α rhythm in the maintenance of sensory information over time.

Second, EEG α in humans is also enhanced in response to certain static stimuli (“flickering wheel illusion”, Sokoliuk and VanRullen, 2013). In that experiment, a static wheel (with 32 spokes) produced an illusory impression of flicker in its center when observed in the visual periphery. Correspondingly, during this illusion the occipital α rhythm of the EEG was the only oscillation that showed a clear response peak. The peak was much decreased in a control condition that did not evoke an illusion of flicker (8-spokes wheel). This study showed that alpha oscillations can reverberate in response to steady visual stimulation, and suggested that these reverberations can sometimes exceed the perceptual threshold and be directly experienced as an illusory flicker. Further, this α reverberation appeared to depend on the spatial content of the stimulus.

Together, these two studies using EEG in humans have shown that α is not only a spontaneous oscillation but also reflects visual input processing. What is the cortical spatial profile for this stimulus-related α rhythm? Human EEG cannot easily answer this question due to its poor spatial resolution. Here, we propose to use voltage-sensitive dye imaging (VSD) in V1 of three anesthetized monkeys to investigate at high spatial and temporal resolutions whether the same visual stimulus patterns, perceptual echoes and flickering wheel (see Figure 1), could induce an oscillatory response in Monkey V1 at the population mesoscopic level. Similar to human “perceptual echoes”, we observed a ~10hz spectral peak in the cross-correlation between the stimulus luminance sequence and the corresponding VSD response on each trial. Similarly, a 8-spokes static wheel also elicited a 10hz oscillation in the VSD response, as in the “flickering wheel” illusion.

**Figure 1:**
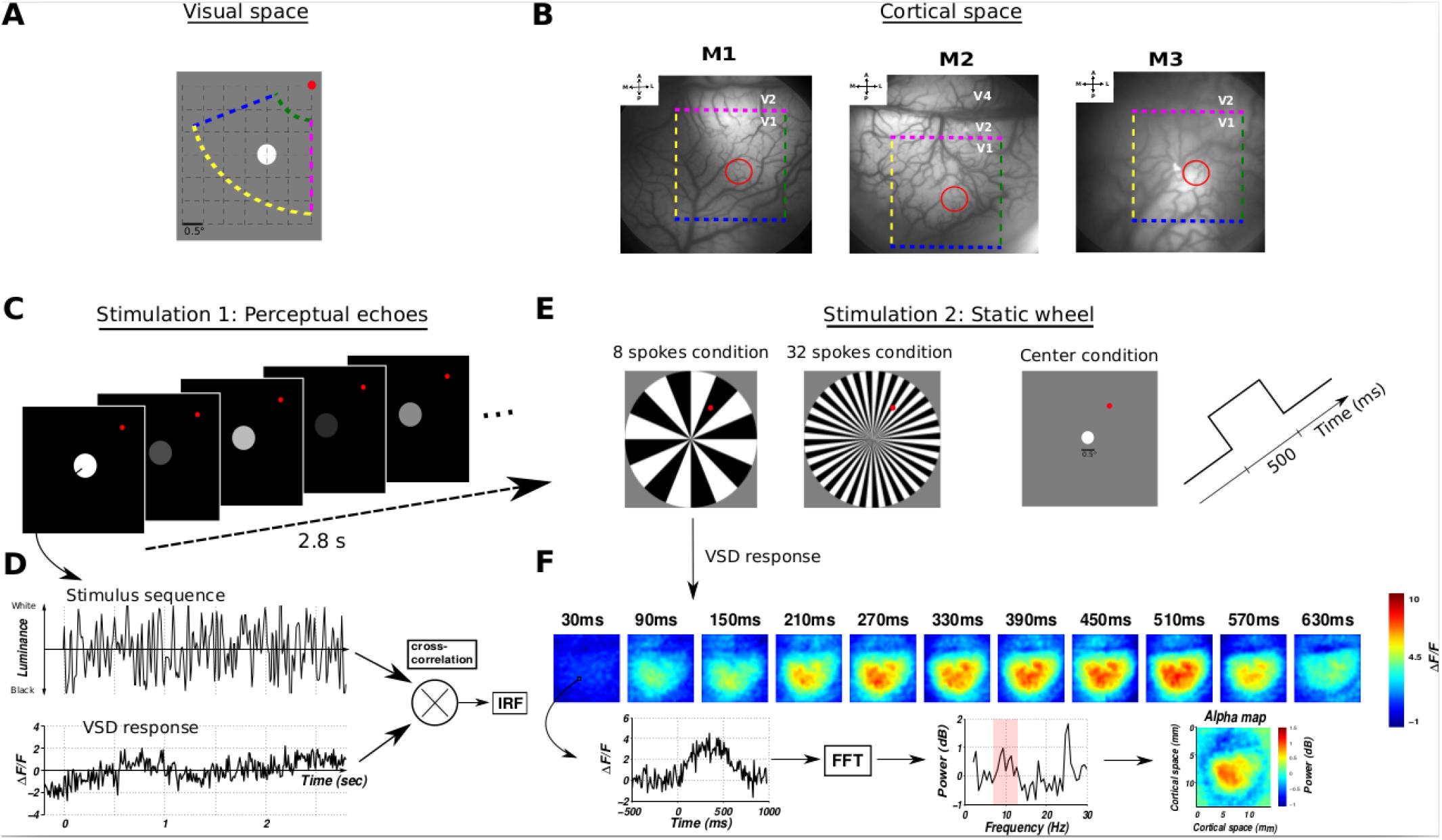
General methodology. **A:** Visual stimuli are presented to three anesthetized monkeys in their bottom left visual field, while recorded using VSDI in their right visual cortex. The red dot indicates the position of the fovea. **B:** Corresponding retinotopic representations in the three cortical spaces, respectively for monkeys M1, M2 and M3. The borders V1/V2 (pink dotted lines) were determined based on the ocular dominance maps (see Supplementary S1). Monkeys were visually stimulated with two protocols. **C:** The “perceptual echoes” stimulation consisted in the presentation of white-noise luminance sequences within a disc stimulus during 2.8 s (with pre-stimulus delay of 200 ms). Figure not drawn to scale. **D:** The cross-correlation analysis between each stimulus sequence and the corresponding VSD response (both were normalized using z-scores) provided an estimate of the impulse response function (IRF) of the visual system. **E:** The “static wheel” stimulation consisted in the presentation of stationary wheels, either composed of 8 spokes or 32 spokes, during 500 ms (with pre- and post-stimulus delays of 500 ms). A 1° diameter Gaussian blob condition presented at the center position of the wheel stimuli was also used as a control. Figure not drawn to scale. **F:** The spectral analysis of the spatiotemporal VSD responses corresponding to each condition produced power spectra in low frequency range, giving the possibility to explore α-band oscillations in space (spatial α maps).

## Results

α-band oscillations (centered around 10 Hz) have been extensively studied in the human visual system using scalp-recorded EEG (Klimesch et al., 1998; Romei et al., 2008), whereas in non-human primates, studies are sparse (Gilad et al., 2012) and stem from local field potentials (LFP) recordings (Wilke et al., 2006; Spaak et al., 2012). To investigate the existence and properties of these oscillations in monkey V1, we proposed a mesoscopic approach using VSD recordings from V1 of three anesthetized monkeys (Fig. 1A-B), visually stimulated with either random sequences of luminance values within a disc stimulus (« perceptual echoes », Fig. 1C-D) or static images of wheels retinotopically centered on the retinotopic representation of the recorded cortical region (« static wheel », Fig. 1E-F). These two stimulation protocols have already demonstrated an enhanced α oscillatory response in humans (VanRullen and Macdonald, 2012; Sokoliuk and VanRullen, 2013). Voltage-sensitive dye imaging signals reflect with a high spatiotemporal resolution the dynamics of cortical processing at the population level, i.e. each pixel represents the sum of membrane potential changes of about 150-200 neurons (Chemla and Chavane, 2010b). Therefore, VSDI provides us with the possibility of exploring the emergence of visually-evoked α oscillations in V1 and at high spatial resolution.

### Visual stimuli can generate enhanced α oscillatory responses in monkey V1

First, we looked at the cross-correlation between dynamic sequences randomly modulated in luminance (i.e. « perceptual echoes » stimulation protocol, see Fig. 1C-D) and the corresponding VSD response. Figure 2A shows the averaged cross-correlation functions across trials (and across the entire V1 cortical region-of-interest) for the three monkeys, respectively M1, M2 and M3. As in humans (VanRullen & Macdonald, 2012), we can observe for each monkey an early-evoked response (marked by a solid arrow) followed (dashed arrows in M1 and M2) or not (M3) by an α oscillation. This oscillation was further characterized in Figure 2B by exploring the power spectrum of the cross-correlation computed pixel-by-pixel and averaged (black traces). In order to verify the stimulus-related nature of this ~10 Hz oscillation, we also performed the same analysis on the cross-correlation functions between each input sequence and the VSD response from a randomly chosen trial. As expected, the averaged power spectrum of this surrogate, depicted in red on each panel of Figure 2B, is lacking the α peak. Figure 2C shows the z-score analysis between these two conditions (the surrogate being averaged over 100 repetitions) for each frequency. This analysis shows that a stimulus-induced increase in oscillation occurs only at ~10 Hz only for M1 and M2 (threshold of 3.09 z-score which corresponds to p-value of 0.01).

**Figure 2:**
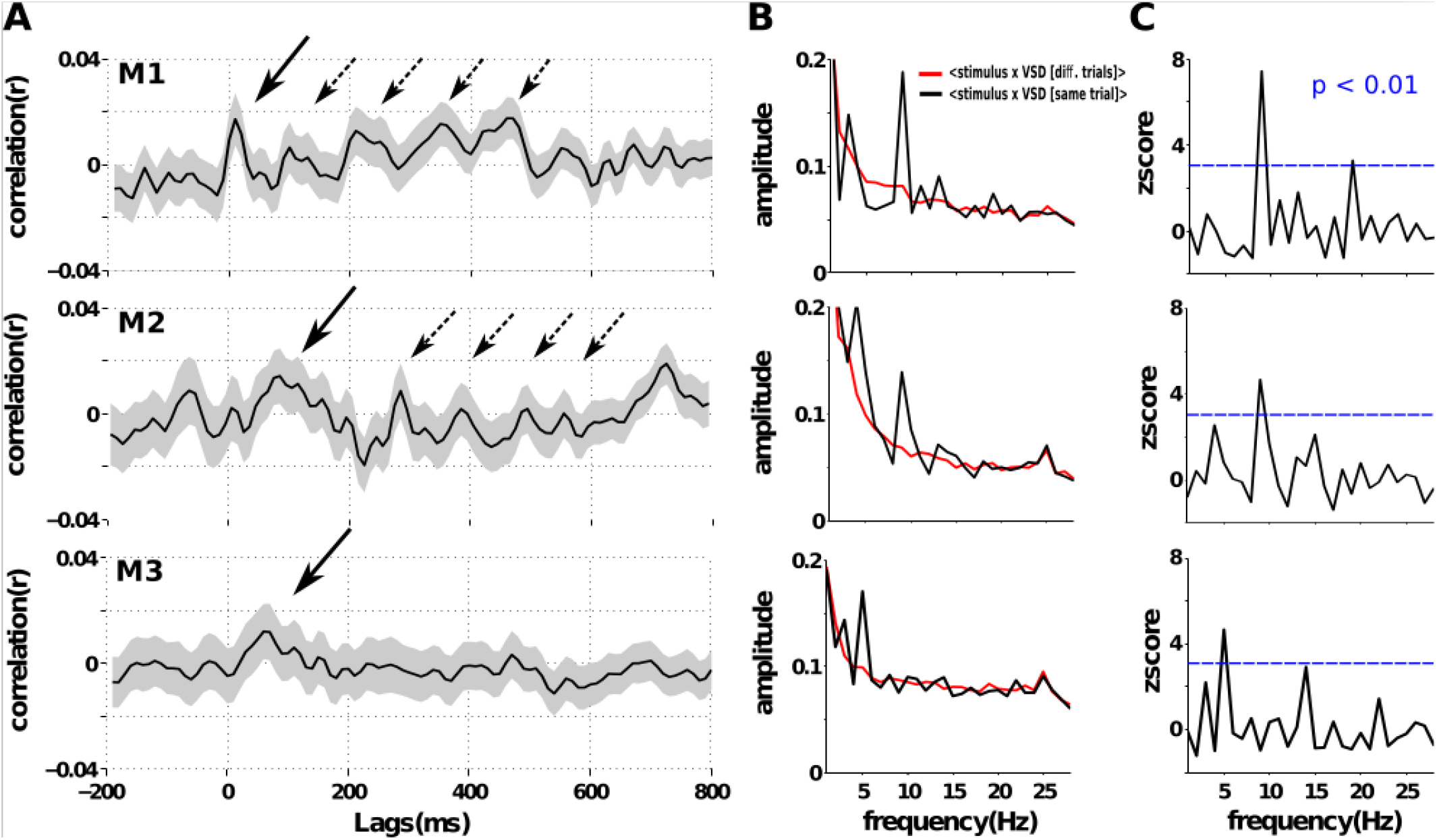
“Perceptual echoes” protocol. **A:** Impulse response functions (see Fig. 1C-D for the methodology) averaged over trials (shaded areas represent SEM across trials) for monkey M1, M2 and M3. A visual evoked response was present in the three monkeys (solid arrows), whereas only monkeys M1 and M2 exhibited an oscillatory response (dashed arrows). **B:** Corresponding amplitude spectra of the cross-correlation functions (black traces) for the three monkeys. The same analysis was also done on the cross-correlation functions between each input sequence and the VSD response from a randomly chosen trial (red traces). **C:** Z-score analysis between the black and red curves (averaged over 100 repetitions) shown in B. A threshold of 3.09 z-score (horizontal blue lines) corresponds to a p-value of 0.01 and considered statistically significant.

Second, we looked at the spectral analysis of VSD responses to static wheel stimuli (i.e. « static wheel » stimulation protocol, see Fig. 1E-F) presented with various spatial frequencies. Time-courses of the VSD responses for each stimulation condition (Gaussian stimulus, 8- and 32-spokes wheel stimuli represented in black, blue and red respectively) are plotted for the three monkeys in Figure 3, respectively in panels **A**, **D** and **G.** The corresponding power spectra, computed by Fourier transformation of the VSD data, are then analyzed. One static wheel stimulus (8 spokes) evoked a clear ~10 Hz oscillatory peak (Figure 3, blue power spectrum for 8-spokes wheel stimuli) for the same two monkeys M1 (Fig. 3B-C) and M2 (Fig. 3E-F), but not M3 (Fig. 3H-I). Stars on Fourier plots denote a statistically significant increase of power magnitude relative to a 1/f power spectrum (nonparametric one-tailed Wilcoxon signed rank test, p < 0.001) computed for each frequency in the α range (6-11 Hz for M1 and 7-12Hz otherwise) of the power spectra.

**Figure 3:**
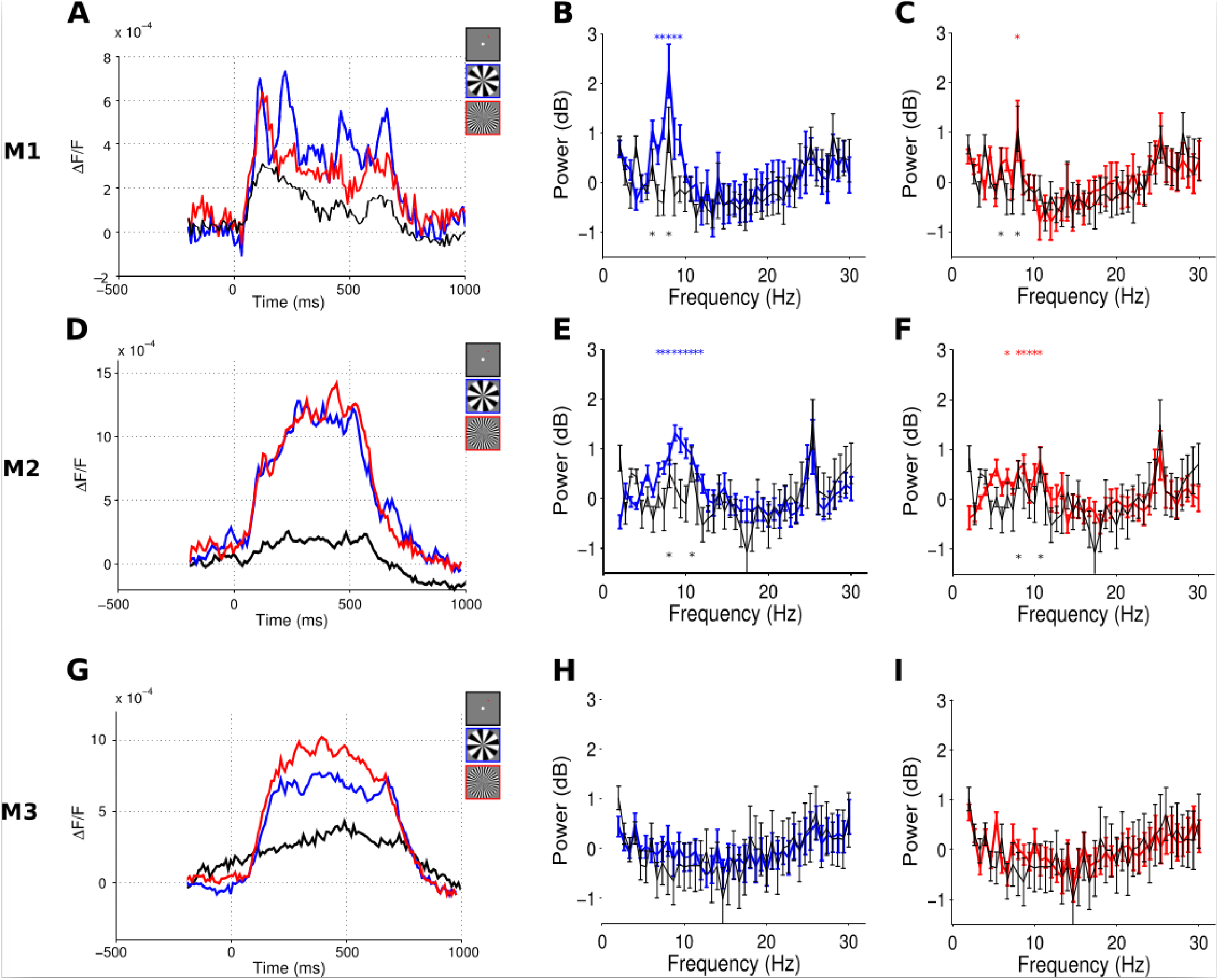
“Static wheel” protocol. The time-courses of the VSD responses for the three stimulation conditions (see Fig. 1E-F for the methodology; Gaussian stimulus, 8- and 32-spokes wheel stimuli represented in black, blue and red respectively) are plotted for the three monkeys in panels **A**, **D** and **G** respectively. The corresponding power spectra, computed by Fourier transformation of the VSD data, are plotted in **B** (8-spokes wheel) and **C** (32-spokes wheel) for M1, **E** and **F** for M2 and **H** and **I** for M3. The power spectrum of the Gaussian stimulation is superimposed in each graph in black. Stars denote a statistically significant increase of power magnitude relative to a 1/f power spectrum (nonparametric one-tailed Wilcoxon signed rank test, p < 0.001) computed for each frequency in the α range (gray-shaded area) of each power spectra.

### α oscillatory activity in monkey V1 depends on the spatial frequency of the stimulus

We then wondered whether this power increase in the α-band systematically varies with the visual stimuli. For both monkeys, the α peak seems much stronger for the 8-spokes wheel condition (blue traces) than for the 32-spokes wheel condition (red traces). To test this observation, we combined the two wheel conditions (8- and 32-spokes) and extracted the peak frequency of the resulting power spectrum (8 Hz for M1 and 8.66 Hz for M2) and we derived the spatial alpha maps separately for all conditions at that peak frequency (see Figure 4, *bottom rows* for M1 and M2: Gaussian stimulus, 8- and 32-spokes wheel stimuli in panels A, B and C respectively). In panel D of Figure 4, we reported the averaged α amplitude over a specific ROI, shown as black contours on top of all maps. This ROI corresponds to the cortical representation of the wheel center (highest-level activity pixels in V1 in response to the local stimulation shown in panel A). For both M1 and M2, the averaged alpha amplitude was significantly stronger for the 8-spokes wheel stimulus (in blue) than for the other two conditions, i.e. the local stimulus (in black) and the 32-spokes wheel stimulus (in red; nonparametric Wilcoxon rank sum test, p < 0.001).

**Figure 4:**
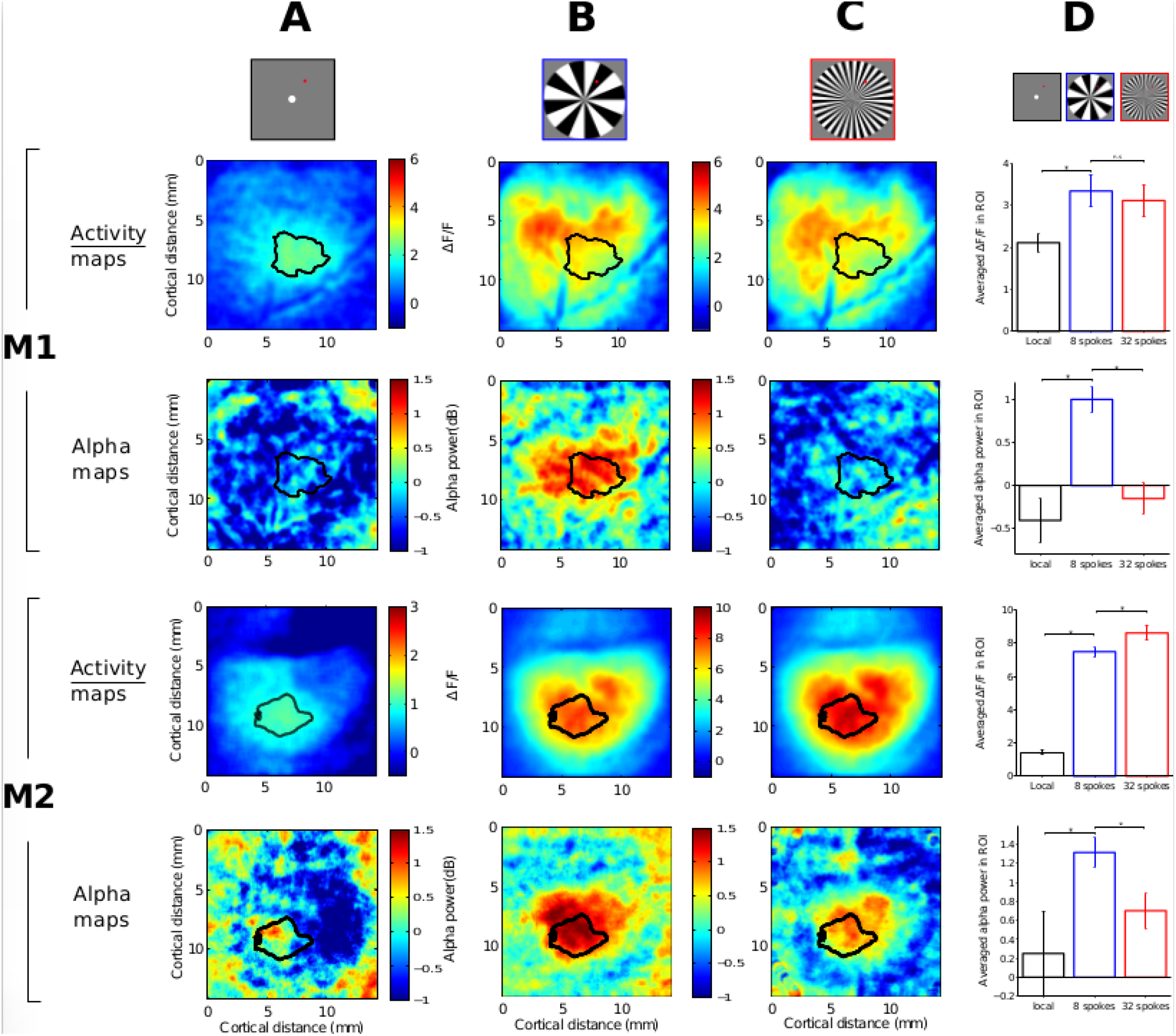
Spatial activity maps (*top rows*) and spatial α maps (*bottom rows*) for monkeys M1 and M2 in response to **A:** the local Gaussian stimulus, **B:** the 8-spokes wheel condition, **C:** the 32-spokes wheel condition. **D:** Averaged VSD activity (*top rows*) and averaged α activity (*bottow rows*) over the ROI shown as black contours on top of all maps to quantitatively compare the three visual stimulation conditions (Local, 8- and 32-spokes wheels in black, blue and red respectively). * p<0.001, n.s non-significant (Wilcoxon rank sum test).

### α oscillatory activity in monkey V1 can be dissociated from evoked activity

Qualitatively, both wheel stimuli generated very similar VSD responses (see Fig. 3, A-D-G for the three monkeys respectively), i.e. in amplitude and time-course, however only the 8 spokes evoked a strong alpha activity. This suggests that there was no trivial link between alpha oscillatory activity and overall evoked activity level. A quantitative analysis is reported in Figure 4D (*top rows* for M1 and M2), by temporally averaging the VSD activity over the steady-state period (spatial activity maps in panels A, B and C, *top rows*). We observed a small difference in VSD activity for M1 between the two wheel conditions (8- vs. 32-spokes wheel; nonparametric Wilcoxon rank sum test, p = 0.0143) and a significant increase in VSD activity for M2 in response to the 32-spokes wheel stimulus compared to the 8-spokes wheel stimulus (nonparametric Wilcoxon rank sum test, p < 0.001). In both cases, these results show that there is no link between α oscillatory activity and the global level of VSD evoked activity.

### α oscillatory activity in monkey V1 is maximal at the wheel’s center and decreases rapidly with eccentricity

Finally, we explored the spatial dependence of the α activity relative to the center of the wheel stimuli (specifically in the 8-spokes wheel condition, where the strongest alpha response was recorded). For both monkeys, we determined the center of mass of the local ROI depicted in black in Figure 4, and from this point we computed an isotropic distance map (see Figure 5A) in order to plot both the VSD activity (in black) and the α amplitude (in blue) distributions in response to the 8-spokes wheel stimulus (see respective maps in Fig. 4B) as a function of eccentricity from center. The distributions were normalized to their maximum and fitted to a 1D Gaussian function of the form:

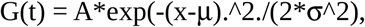

**Figure 5:**
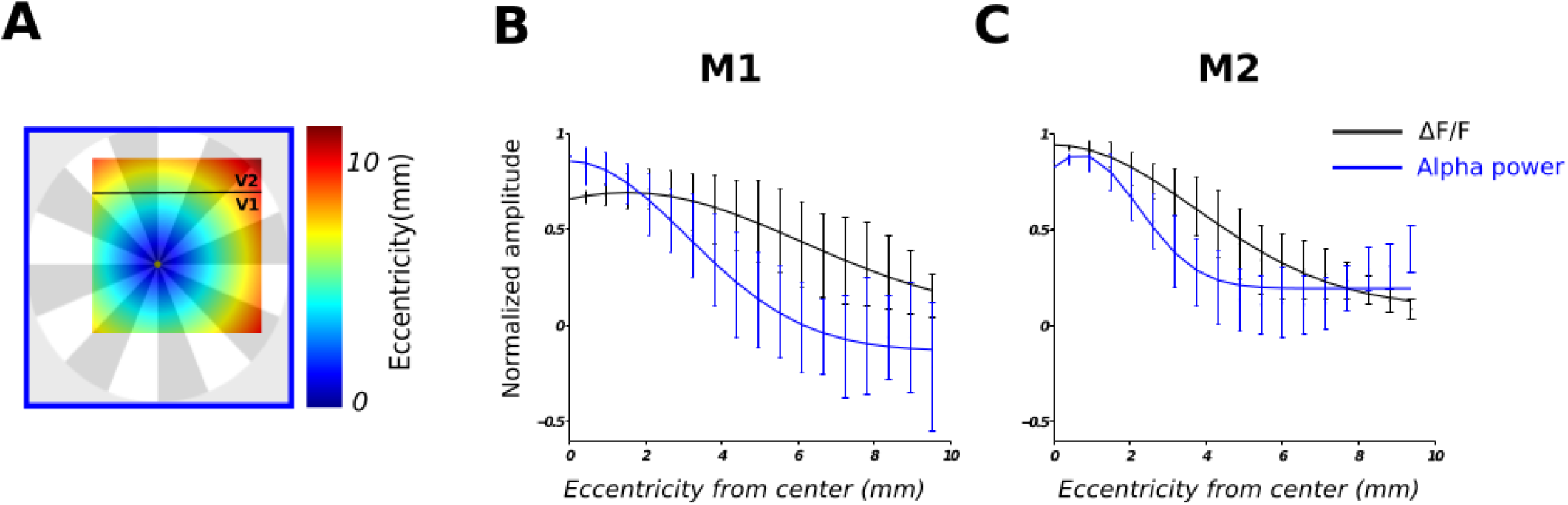
Spatial dependence of α and VSD activity relative to the center of the 8-spokes wheel condition. **A:** Isotropic distance map from the center position of the wheel and limited to pixels in the V1 region. The center positions for M1 and M2 were computed as the centers of mass of the local ROIs depicted in black in Figure 4, for M1 and M2 respectively. The color code represents the cortical eccentricity from this point expressed in millimeters. After normalization to their maximum, the distributions (mean ± SD) of VSD amplitude (black lines) and α amplitude (blue lines) in response to the 8-spokes wheel stimulus (see maps in Fig. 4B) were plotted as a function of eccentricity from center in panel **B** for M1 and in panel **C** for M2.

where μ and σ are respectively the mean and the standard deviation. The amplitude distributions (mean ± SD) and fits were then superimposed in Figure 5B for M1 and Figure 5C for M2. This analysis provided us with two interesting results: i) the strongest α oscillatory activity lies in or around the retinotopic position of the wheel’s center (blue curves, μ = 0 and μ = 0.4 for M1 and M2 respectively), consistent with subjective reports of the flickering wheel illusion in human subjects; Sokoliuk and VanRullen, 2013); ii) α oscillatory activity is more local since it decreases rapidly along cortical space (blue curves, σ = 3.1 and σ = 2.0 for M1 and M2 respectively), whereas the VSD response extends much further (black curves, σ = 4.6 and σ = 4.0 for M1 and M2 respectively). Note that another recording session in monkey M2 one week later showed reproducibility of the results (see Supplementary Figure).

## Discussion

We used voltage-sensitive dye imaging from V1 in three anesthetized monkeys to investigate the presence of oscillatory activity in response to specific visual stimulation previously tested in human EEG. In two monkeys, we observed a ~10hz spectral peak in the cross-correlation between random, non-periodic dynamic luminance sequences and the corresponding VSD response on each trial. The same two monkeys also showed a ~10hz oscillatory response when visually stimulated with a static wheel. In conclusion, both visual stimulation protocols (i.e. « perceptual echoes » and « static wheel », see methods) induced α-band oscillations in V1 of two of the three monkeys. Altogether, our results are in accordance with the human studies (respectively VanRullen and MacDonald, 2012 and Sokoliuk and VanRullen, 2013): α-frequency reverberations can be observed in response to specific visual stimuli; the magnitude of this alpha response depends on the type of stimulation (spatial frequency of the wheel), but it can be dissociated from stimulus-evoked activity. Therefore the current study demonstrates that, in monkeys, α-band activity, generally thought of as spontaneous fluctuations or an ‘idling rhythm’, can also be linked to visual processing.

Interestingly, not all monkeys exhibited α oscillations. This inter-individual variability in monkeys is reminiscent of inter-individual differences in human α rhythm. The latter have been widely studied (Klimesch et al., 1999; 2003; Olbrich and Achermann, 2005; Bodenmann et al., 2009) and have been shown to be large and mainly dependent on age and genetic factors. For example, in a recent study dedicated to inter- and intra-individual variability in alpha peak frequency, Haegens and collaborators reported a lack of alpha peak in the spectra of some participants (Haegens et al., 2014). However, the reason for such notable difference in those individuals remains unanswered.

Inspecting the α dependence on the spatial frequency of the wheel stimulus (8- vs 32-spokes wheel), we surprisingly reported that humans and monkeys show opposite preference. Indeed, the spatial frequency of the wheel pattern that generated the strongest α fluctuations in monkeys (8-spokes; Fig. 4B) does not induce an oscillatory perception in humans (Sokoliuk and VanRullen, 2013; their Fig 2A). Although an hypothesis about difference in consciousness states cannot be totally rejected (Purdon et al., 2013), we rather suggest that the relative preference is more likely to be explained by differences in the spatial organization of the visual cortex between the two species (Dow et al., 1981; Van Essen et al., 1984; Dumoulin and Wandell, 2008; Harvey and Dumoulin, 2011), e.g. cortical magnification factor, receptive field size, eccentricity maps, and remains to be explored.

The current study analyzing VSDI recordings in monkey V1 gave us the opportunity to explore the neural basis of stimulus-related α oscillations. This question is indeed difficult to address, since α-band oscillations are ubiquitous in the brain. They have been simultaneously observed in many brain regions, the thalamus and the cortex being the two major candidate generators. The debate on the genesis of α-band oscillations mainly focused on thalamocortical resonance (Steriade et al., 1990; Bollimunta et al., 2011) and cortical feedback (Buffalo et al., 2011; Van Kerkoerle et al., 2011) mechanisms. However, finding the origins of alpha-frequency oscillations does not necessarily reveal whether they play a role in visual information processing. Here, we showed that an oscillatory activity in the α-band occurs in V1 and that this oscillation can be induced by the visual stimulation. Therefore, we explicitly provided a link between the α rhythm and visual input processing in monkey V1. Furthermore, we revealed that the α peak is localized in space at the retinotopic position of the wheel’s center while it is not necessarily the case for the peak of VSD activity; and the spatial extent of alpha reverberations was also limited compared to the corresponding VSD evoked activity (Fig. 5). This localized reverberatory activity could therefore be part of the neural substrates accounting for the perception of an illusory regular flicker restricted to the wheel center, previously reported in humans (Sokoliuk and VanRullen, 2013).

Finally, this study has been done on anesthetized monkeys in order to demonstrate that α oscillations are stimulus-induced and not resulting from task-dependent feedback signals related to attention (van Kerkoerle et al., 2014) or other higher-order phenomenon (Gilbert and Li, 2013). In addition, with anesthetized and paralyzed monkeys, we ruled out the possible idea that the α rhythm could have been solely induced by small microsaccades made during fixation (Dimigen et al., 2011). We can wonder whether the same oscillations could be induced in awake monkeys. Indeed, ongoing activity in the α-band is well known to be prominent in a quiet awake state and linked to conscious perception (Ruhnau et al., 2014) and attention (Klimesch 2012), all of which are likely to be negatively affected by anaesthesia. However, as we focused our study on stimulus-related rather than spontaneous α activity, we surmise that our results should not solely reflect the anesthesia level (although anesthesia could be responsible for other mechanisms in the α-band (e.g. Vijayan et al., 2013; Flores et al., 2017)). Nevertheless, testing awake monkeys with behavioral feedback will be a straightforward follow-up to the present study, to confirm that monkeys perceive the stimuli as humans do, and deepen our knowledge, still largely incomplete, about the neural basis of α-band oscillations.

## Materials and Methods

### Experimental procedures

The experiments were conducted on three monkeys (three female macaca fascicularis). Experimental protocols have been approved by the Marseille Ethical Committee in Neuroscience (approval A10/01/13, official national registration 71-French Ministry of Research). All procedures complied with the French and European regulations for animal research, as well as the guidelines from the Society for Neuroscience.

### Surgical preparation, anesthesia and VSD imaging protocol

The monkeys were chronically implanted with a recording chamber over the right hemisphere. Subsequently the dura-mater of the primary visual cortex (V1) was removed surgically over the recording aperture (18mm diameter) and a silicon-made artificial dura-mater was inserted under aseptic conditions to insure a good preparation and an optical access to the cortex (Arieli et al., 2002). The day of a recording session, the monkeys were initially anesthetized with intramuscular Ketamine (10mg/kg), Xylazine (0.5mg/kg) and Robinul (0.1mg/kg). Then continuous anesthesia was induced with an intravenous (IV) propofol bolus (2.5mg/kg) and maintained with slow IV propofol infusion (0.3mg/kg/min), while monkeys were artificially ventilated. Animals were then paralyzed with rocuronium bromide (esmeron, 0.5mg/kg/h), the left eye was dilated using atropine and a contact lens was placed to prevent drying. During the experimental protocol, physiological parameters, i.e. temperature, heart frequency, concentration of expired carbon dioxide and oxygen saturation level, were monitored every 30 minutes. At the end of experimental recordings, propofol and esmeron infusions were stopped and the animal recovered in a very short time (less than 20 minutes on average). The train-of-four (TOF) twitch technique was used to verify induction and reversal of paralysis (Hughes and Griffiths, 2002). Before recordings, the cortex was stained for three hours with the voltage-senstive dye RH-1691 (Optical Imaging) prepared in artificial cerebrospinal fluid (aCSF) at a concentration of 0.2 mg/ml and filtered through a 0.2 μm filter. After this staining period, the chamber was rinsed thoroughly with filtered aCSF to wash out any supernatant dye, the artificial dura-mater was inserted back in position and the chamber was closed with transparent agar and cover glass. Subsequently, optical signals were recorded from a focal plane ~300 μm below the cortical surface using a Dalstar camera (512 × 512 pixels resolution, frame rate of 110 Hz) driven by the Imager 3001 system (Optical Imaging). Excitation light was provided by a 100W halogen lamp filtered at 630 nm and fluorescent signals were high-pass filtered at 665 nm. The surgical preparation and VSD imaging protocol have been described in detail in Reynaud et al. (2012).

### Visual stimulation

`The` visual stimuli consisted of either randomly generated sequences (2.8 sec duration, 80 trials) of luminance values (from 0 to 255 screen gun values, from 2 to 130 cd.m-^2^) displayed within a peripheral disc stimulus (radius: 1.5 degrees of visual angle) on a black background after a pre-stimulus period of 200 ms (“perceptual echoes” protocol) or png images of 8- and 32-spokes wheels (radius: 13.77 degrees of visual angle, 30 trials) displayed on a gray background for 500 ms after a pre-stimulus period of 500 ms (“static wheel” protocol). In addition, a blank condition, i.e. where no visual stimulus was presented on the screen, was also used in both protocols for the corresponding duration and same number of trials, whereas a local stimulation condition, i.e. 1° diameter Gaussian blob, was only used for the static wheel protocol. Stimuli were displayed monocularly at 60 Hz on a gamma corrected LCD monitor (placed at 57 cm distance from the animal eye plane) using the Elphy software (G. Sadoc, Unic, Paris), communicating with the VDAQ acquisition system (Optical Imaging). The position of the fovea was initially located with a retinal angiography, projected back on the screen using a mirror and all stimuli were centered on the bottom-left at (−1°, −2°) from the foveal position (red dots in Fig. 1A-C-E, not visible during recordings).

### Data analyses

Stacks of images were stored on hard-drives for off-line analysis with MATLAB R2014a (MathWorks), using the Optimization, Statistics, and Signal Processing Toolboxes.

### VSD evoked activity

For both protocols, the evoked response to each stimulus was computed in three successive basic steps. First, the recorded value at each pixel was divided by the average value before stimulus onset (frames 0 division) to remove slow stimulus-independent fluctuations in illumination and background fluorescence levels. Second, this value was subsequently subtracted by the value obtained for the blank condition (blank subtraction) to eliminate most of the noise due to heartbeat and respiration (Shoham et al., 1999). Third, a linear detrending of the timeseries was applied to remove residual slow drifts induced by dye bleaching (Chen et al., 2008; Meirovithz et al., 2010). In addition, for the “static wheel” protocol, a spatial smoothing (convolving the raw matrix with a 5×5 pixel flat matrix) and spatial z-score normalization were applied on each evoked averaged map (Fig. 4, *top rows*). The latter was done by dividing them by the standard deviation of the blank condition map, pixel by pixel. In Figure 3, the time-course traces (*top row*) result from a spatial averaging over a specific region of interest (ROI), shown as black contours in Figure 4. These ROIs correspond to high-level activity pixels in V1 in response to the local stimulation, i.e. stimulation of the center of the wheel stimuli (Fig. 4A, z-score = 2).

### Frequency analysis

For the “perceptual echoes” protocol, we first estimated pixel-by-pixel the cross-correlation function between each VSD trial and the corresponding visual stimulation sequence resampled at 100 Hz. For each pixel, we averaged the cross-correlation functions across trials (after excluded the first 0.5 s of each VSD trial and each stimulus sequence) and then performed a Fourier transform of this averaged cross-correlation function to derive its amplitude spectrum. Finally, we computed the averaged power spectrum across pixels (over the entire V1 cortical region-of-interest). For the “static wheel” protocol, power spectra for each condition were directly computed by Fourier transformation of the single-trial VSD data, pixel-by-pixel, and averaged across trials. Corresponding 1/f fits computed over the range of frequencies from 2 to 30 Hz were subsequently removed to obtain the final amplitude spectra. Spatial α maps can then be derived by specifically looking at one (e.g. peak frequency of the spectrum) or the average of several frequencies in the α range (6-11 Hz for M1 and 7-12Hz otherwise).

### Statistical procedure

We used two-tailed nonparametric Wilcoxon signed rank test for two-sample or paired data with P<0.001 considered significant, except where otherwise stated (one-tailed).

## Acknowledgments

The authors are thankful to Pierre Simeone, and the CE2F-PRIM team, for their help with the experiments. The authors acknowledge fundings from the ERC Consolidator grant P-CYCLES number 614244, the ANR BalaV1 (ANR-13-BSV4-0014-02) and FRQS Vision Health Research Network of Quebec networking grant.

## Competing interests

The authors declare no competing interests.

---

**Supplementary Figure:**
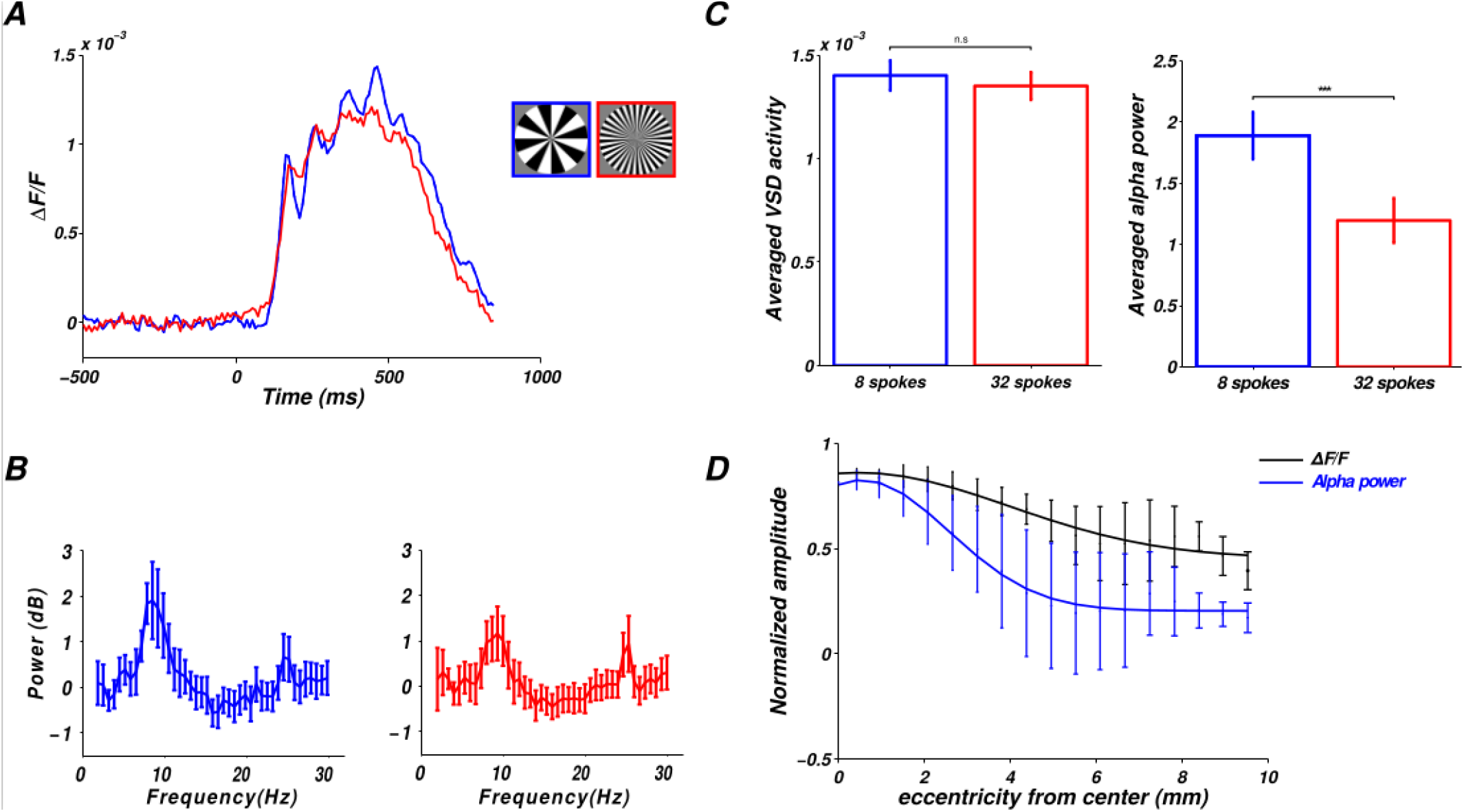
Results reproducibility. Another recording session in monkey M2 one week later reported a strong α power increase in response to the 8-spoke wheel stimulus (blue traces) with the same characteristics: its dissociation with VSD evoked activity (**A-C**), its dependence with the spatial frequency of the stimulus (**B-C**) and its spatial relationship with the center of the wheel (**D**).

## Notes

+ Co-supervisors

*Conflict of Interest*: The authors declare no competing interests.

#### Summary of Updates

Reference in the methods section revised

## References

Arieli, A., Grinvald, A., & Slovin, H. (2002). Dural substitute for long-term imaging of cortical activity in behaving monkeys and its clinical implications. Journal of neuroscience methods, 114(2), 119–133.

Berger, H. (1929). Über das elektrenkephalogramm des menschen. European Archives of Psychiatry and Clinical Neuroscience, 87(1), 527–570.

Bodenmann, S., Rusterholz, T., Dürr, R., Stoll, C., Bachmann, V., Geissler, E., … & Landolt, H. P. (2009). The functional Val158Met polymorphism of COMT predicts interindividual differences in brain α oscillations in young men. Journal of Neuroscience, 29(35), 10855–10862.

Bollimunta, A., Mo, J., Schroeder, C. E., & Ding, M. (2011). Neuronal mechanisms and attentional modulation of corticothalamic alpha oscillations. Journal of Neuroscience, 31(13), 4935–4943.

Buffalo, E. A., Fries, P., Landman, R., Buschman, T. J., & Desimone, R. (2011). Laminar differences in gamma and alpha coherence in the ventral stream. Proceedings of the National Academy of Sciences, 108(27), 11262–11267.

Busch, N. A., Dubois, J., & VanRullen, R. (2009). The phase of ongoing EEG oscillations predicts visual perception. Journal of Neuroscience, 29(24), 7869–7876.

Chemla S, Chavane F (2010). Voltage-sensitive dye imaging: Technique review and models. J Physiol (Paris) 104:40–50.

Chemla, S., & Chavane, F. (2010). A biophysical cortical column model to study the multi-component origin of the VSDI signal. Neuroimage, 53(2), 420–438.

Chen Y, Geisler WS, Seidemann E (2008). Optimal temporal decoding of neural population responses in a reaction-time visual detection task. Journal of neurophysiology, 99:1366–1379.

Dow, B. M., Snyder, A. Z., Vautin, R. G., & Bauer, R. (1981). Magnification factor and receptive field size in foveal striate cortex of the monkey. Experimental Brain Research Experimentelle Hirnforschung Expérimentation Cérébrale, 44(2), 213–228.

Dimigen O, Werkle-Bergner M, Meyberg S, Kliegl R and Sommer W (2011). Microsaccades and EEG alpha oscillations: a close relationship? Front. Hum. Neurosci. Conference Abstract: XI International Conference on Cognitive Neuroscience (ICON XI). doi: 10.3389/conf.fnhum.2011.207.00128

Dumoulin, S. O., & Wandell, B. A. (2008). Population receptive field estimates in human visual cortex. NeuroImage, 39(2), 647–660.

Eckhorn, R., Frien, A., Bauer, R., Woelbern, T., & Kehr, H. (1993). High frequency (60-90 Hz) oscillations in primary visual cortex of awake monkey. Neuroreport, 4(3), 243–246.

Flores, F. J., Hartnack, K. E., Fath, A. B., Kim, S. E., Wilson, M. A., Brown, E. N., & Purdon, P. L. (2017). Thalamocortical synchronization during induction and emergence from propofol-induced unconsciousness. Proceedings of the National Academy of Sciences, 114(32), E6660–E6668.

Gilad, A., Meirovithz, E., Leshem, A., Arieli, A., & Slovin, H. (2012). Collinear stimuli induce local and cross-areal coherence in the visual cortex of behaving monkeys. PloS one, 7(11), e49391.

Gilbert, C. D., & Li, W. (2013). Top-down influences on visual processing. Nature Reviews Neuroscience, 14(5), 350–363.

Haegens, S., Cousijn, H., Wallis, G., Harrison, P. J., & Nobre, A. C. (2014). Inter-and intra-individual variability in alpha peak frequency. Neuroimage, 92, 46–55.

Hanslmayr, S., Aslan, A., Staudigl, T., Klimesch, W., Herrmann, C. S., & Bäuml, K. H. (2007). Prestimulus oscillations predict visual perception performance between and within subjects. Neuroimage, 37(4), 1465–1473.

Harvey, B. M., & Dumoulin, S. O. (2011). The Relationship between Cortical Magnification Factor and Population Receptive Field Size in Human Visual Cortex: Constancies in Cortical Architecture. Journal of Neuroscience, 31(38), 13604–13612.

Hughes, S., & Griffiths, R. (2002). Anaesthesia monitoring techniques. Anaesthesia and Intensive Care Medicine. The Medicine Publishing Company Ltd, pp. 477–480.

Jensen, O., & Mazaheri, A. (2010). Shaping functional architecture by oscillatory alpha activity: gating by inhibition. Frontiers in human neuroscience, 4, 186.

Klimesch, W., Doppelmayr, M., Russegger, H., Pachinger, T., & Schwaiger, J. (1998). Induced alpha band power changes in the human EEG and attention. Neuroscience letters, 244(2), 73–76.

Klimesch, W. (1999). EEG alpha and theta oscillations reflect cognitive and memory performance: a review and analysis. Brain research reviews, 29(2), 169–195.

Klimesch, W., Sauseng, P., & Gerloff, C. (2003). Enhancing cognitive performance with repetitive transcranial magnetic stimulation at human individual alpha frequency. European Journal of Neuroscience, 17(5), 1129–1133.

Klimesch, W., Sauseng, P., & Hanslmayr, S. (2007). EEG alpha oscillations: the inhibition–timing hypothesis. Brain research reviews, 53(1), 63–88.

Klimesch, W. (2012). Alpha-band oscillations, attention, and controlled access to stored information. Trends in cognitive sciences, 16(12), 606–617.

Lange, J., Keil, J., Schnitzler, A., van Dijk, H., & Weisz, N. (2014). The role of alpha oscillations for illusory perception. Behavioural brain research, 271, 294–301.

Meirovithz E, Ayzenshtat I, Bonneh YS, Itzhack R,Werner-Reiss U, Slovin H (2010) Population response to contextual influences in the primary visual cortex. Cereb Cortex 20:12930–1304.

Milton, A., & Pleydell-Pearce, C. W. (2016). The phase of pre-stimulus alpha oscillations influences the visual perception of stimulus timing. NeuroImage, 133, 53–61.

Muller, L., Reynaud, A., Chavane, F., & Destexhe, A. (2014). The stimulus-evoked population response in visual cortex of awake monkey is a propagating wave. Nature communications,5.

Olbrich, E., & Achermann, P. (2005). Analysis of oscillatory patterns in the human sleep EEG using a novel detection algorithm. Journal of sleep research, 14(4), 337–346.

Pfurtscheller, G., Stancak, A., & Neuper, C. (1996). Event-related synchronization (ERS) in the alpha band–an electrophysiological correlate of cortical idling: a review. International journal of psychophysiology, 24(1), 39–46.

Purdon, P. L., Pierce, E. T., Mukamel, E. A., Prerau, M. J., Walsh, J. L., Wong, K. F. K., … & Ching, S. (2013). Electroencephalogram signatures of loss and recovery of consciousness from propofol. Proceedings of the National Academy of Sciences, 110(12), E1142–E1151.

Reynaud A, Masson GS, Chavane F (2012) Dynamics of local input normalization result from balanced short-and long-range intracortical interactions in area v1. J Neurosci 32:12558–12569.

Romei, V., Brodbeck, V., Michel, C., Amedi, A., Pascual-Leone, A., & Thut, G. (2008). Spontaneous fluctuations in posterior α-band EEG activity reflect variability in excitability of human visual areas. Cerebral cortex, 18(9), 2010–2018.

Ruhnau, P., Hauswald, A., & Weisz, N. (2014). Investigating ongoing brain oscillations and their influence on conscious perception–;network states and the window to consciousness. Frontiers in Psychology, 5 (1230).

Schmiedt, J. T., Maier, A., Fries, P., Saunders, R. C., Leopold, D. A., & Schmid, M. C. (2014). Beta oscillation dynamics in extrastriate cortex after removal of primary visual cortex. Journal of Neuroscience, 34(35), 11857–11864.

Shoham D, Glaser D, Arieli A, Kenet T, Wijnbergeb C, Toledo Y, Hildesheim R, Grinvald A (1999) Imaging cortical dynamics at high spatial and temporal resolution with novel blue voltage-sensitive dyes. Neuron, 24:791–802.

Sokoliuk R, VanRullen R (2013) The flickering wheel illusion: When α rhythms make a static wheel flicker. The Journal of Neuroscience, 33:13498–13504.

Spaak, E., Bonnefond, M., Maier, A., Leopold, D. A., & Jensen, O. (2012). Layer-specific entrainment of gamma-band neural activity by the alpha rhythm in monkey visual cortex. Current Biology, 22(24), 2313–2318.

Thut, G., Nietzel, A., Brandt, S. A., & Pascual-Leone, A. (2006). α-Band electroencephalographic activity over occipital cortex indexes visuospatial attention bias and predicts visual target detection. Journal of Neuroscience, 26(37), 9494–9502.

Van Essen, D. C., Newsome, W. T., & Maunsell, J. H. (1984). The visual field representation in striate cortex of the macaque monkey: asymmetries, anisotropies, and individual variability. Vision Research, 24(5), 429–448

van Kerkoerle, T. J., Self, M., Poort, J., van der Togt, C., & Roelfsema, P. R. (2011). High frequencies flow in the feedforward direction through the different layers of monkey primary visual cortex while low frequencies flow in the recurrent direction. In Soc. Neurosci. Abstr (Vol. 270).

VanRullen R, Macdonald JS (2012) Perceptual echoes at 10 hz in the human brain. Current biology, 22:995–999.

Vijayan, S., Ching, S., Purdon, P. L., Brown, E. N., & Kopell, N. J. (2013, November). Biophysical modeling of alpha rhythms during halothane-induced unconsciousness. In Neural Engineering (NER), 2013 6th International IEEE/EMBS Conference on (pp. 1104–1107). IEEE.

Wilke, M., Logothetis, N. K., & Leopold, D. A. (2006). Local field potential reflects perceptual suppression in monkey visual cortex. Proceedings of the National Academy of Sciences, 103(46), 17507–17512.

